# Semantic richness and density effects on language production: Electrophysiological and behavioral evidence

**DOI:** 10.1101/509000

**Authors:** Milena Rabovsky, Daniel Schad, Rasha Abdel Rahman

**Affiliations:** Department of Psychology, University of Potsdam, Germany; Department of Linguistics, University of Potsdam, Germany; Department of Psychology, Humboldt-Universität zu Berlin, Germany

**Keywords:** picture naming, ERPs, semantic richness, semantic features, lexical competition

## Abstract

Language production ultimately aims to convey meaning. Yet, words differ widely in the richness and density of their semantic representations and these differences impact conceptual and lexical processes during speech planning. Here, we replicate the recent finding that semantic richness, measured as the number of associated semantic features according to semantic feature production norms, facilitates object naming. In contrast, intercorrelational semantic feature density, measured as the degree of intercorrelation of a concept’s features, presumably resulting in the coactivation of closely related concepts, has an inhibitory influence. We replicate the behavioral effects and investigate their relative time course and electrophysiological correlates. Both the facilitatory effect of high semantic richness and the inhibitory influence of high feature density were reflected in an increased posterior positivity starting at about 250 ms, in line with previous reports of posterior positivities in paradigms employing contextual manipulations to induce semantic interference during language production. Furthermore, amplitudes at the same posterior electrode sites were positively correlated with object naming times between about 230 and 380 ms. The observed effects follow naturally from the assumption of conceptual facilitation and simultaneous lexical competition, and are difficult to explain by language production theories dismissing lexical competition.

## 1 Introduction

Language production aims to convey meaning. Yet, concepts differ substantially in the richness of their semantic representations and in the density of the regions they inhabit in semantic space. For instance, words can be associated with relatively many or few semantic features, and they can co-activate relatively many or few related meaning alternatives. It has recently been shown that this variability modulates language production (Rabovsky, Schad, & Abdel Rahman, 2016). Specifically, object naming was facilitated by semantic richness, quantified by the number of semantic features (e.g. mouse – is small, has four legs etc) associated with a concept based on empirical semantic feature production norms (McRae, Cree, Seidenberg, & McNorgan, 2005). On the other hand, naming was inhibited by intercorrelational feature density, a variable derived from the same feature norms. This variable measures the degree of intercorrelation of a concept’s features (see below for details), which presumably results in increased lexical-semantic co-activation. Relatedly, Mirman (2011) as well as Fieder, Wartenburger, and Abdel Rahman (2018) observed inhibitory influences of the number of a concept’s closely related semantic neighbors (but see Bormann, 2011, who found no influence of overall semantic neighborhood size on object naming in unimpaired subjects). Thus, increasing evidence points to a role of item-endogenous semantic attributes in language production. While semantic neighborhood density and related measures are relatively well-investigated, little is known about effects of semantic richness in language production. Moreover, the electrophysiological correlates and temporal dynamics of these effects are currently unclear, and it has not yet been examined how the neural correlates of these effects relate to those of influences of message-exogeneous contextual manipulations, which are much more common in language production research. The current study aims to address these issues.

Apart from the recent evidence discussed above, semantic factors in language production research have mostly been investigated by manipulating the contexts in which identical messages are produced, rather than manipulating message-endogenous attributes. For example, a semantically related distractor word accompanying a to-be named picture in the picture-word interference (PWI) paradigm (e.g., Glaser & Glaser, 1989; La Heij, 1988; Lupker, 1979; Schriefers, Meyer, & Levelt, 1990), a semantically homogeneous sequence of object pictures in the semantic blocking paradigm (Belke, Meyer, & Damian, 2005; Damian, Vigliocco, & Levelt, 2001; Kroll & Stewart, 1994), or the previous naming of objects from the same semantic category giving rise to cumulative semantic interference (Howard, Nickels, Coltheart, & Cole-Virtue, 2006) slows down object naming compared to unrelated distractor and block conditions. These semantic interference effects have been assumed to reflect competitive mechanisms at the level of lexical selection (Levelt, Roelofs, & Meyer, 1999; Starreveld & La Heij, 1995).

Semantic context has also been observed to yield facilitation (e.g., Abdel Rahman & Melinger, 2007; Alario, Segui, & Ferrand, 2000; Costa, Alario, & Caramazza, 2005; La Heij, Dirkx, & Kramer, 1990). Most language production theories share the assumption that semantic contexts can induce facilitative priming of the to be named target at the conceptual (Abdel Rahman & Melinger, 2009b; Costa, et al., 2005) or lexical processing level (Mahon et al., 2007). This should result in faster activation of the target concept and / or its lexical representation.

While language production theories roughly agree on the explanation of facilitative influences of semantic contexts, there is considerable disagreement concerning why interference overrides facilitation in some situations but not in others. Specifically, there is an active debate as to whether lexical selection is a competitive process (Abdel Rahman & Melinger, 2009a, 2009b; Hantsch & Maedebach, 2013; Jescheniak, Matushanskaya, Mädebach, & Müller, 2014; Jescheniak, Schriefers, & Lemhöfer, 2014; Levelt, et al., 1999; Roelofs, 2018; Roelofs, Piai, & Schriefers, 2013a, 2013b; Starreveld, La Heij, & Verdonschot, 2013) or whether interference effects are due to different mechanisms in different paradigms (Costa, et al., 2005; Finkbeiner & Caramazza, 2006; Janssen, 2013; Mahon & Caramazza, 2009; Mahon, Costa, Peterson, Vargas, & Caramazza, 2007; Navarrete & Mahon, 2013; Oppenheim et al., 2010). Specifically, the response exclusion hypothesis (Mahon, et al., 2007) assumes that interference from related distractor words is located post-lexically in the articulators that constitute a bottleneck stage to which distractor words have privileged access. The difficulty to exclude distractor words from production increases with their response relevance. Categorically related words are potentially relevant responses compared to unrelated words that are quickly identified as irrelevant based on their semantic attributes. For blocking paradigms Oppenheim and colleagues (2010) have suggested an implicit learning mechanism. Following the selection of a target word, connections between the co-activated representations at the conceptual and lexical level are readjusted (weakened) in order to make future retrieval of the target more efficient.

In contrast, the swinging lexical network proposal (SLN) as a variant of competitive models assumes that semantic contexts cause simultaneous conceptual priming and lexical competition. The trade-off between conceptual facilitation and lexical competition is assumed to depend on the activation of a strongly interrelated lexical cohort (Abdel Rahman & Melinger, 2009b; 2019; Melinger & Abdel Rahman, 2012). This variant of lexical competition models was proposed in order to account for semantic context effects of opposite polarity in different naming paradimgs (e.g., Alario, et al., 2000; La Heij, et al., 1990). For instance, when target and distractor in the PWI task are from the same semantic category (e.g. dog and cat), they spread converging activation to additional category members through shared semantic features so that a cohort of interrelated lexical representations is co-activated and competes for selection, resulting in one-to-many competition, which induces substantial interference effects that override conceptual priming. In contrast, when target and distractor are associatively related (e.g. bee and honey), their activation does not converge on semantic features that are shared by further concepts so that they do not jointly activate other related concepts. Instead, in this case both the target and the distractor separately activate other mutually unrelated concepts so that their activation diverges and eventually dissipates. Thus, only the target and distractor are highly activated, resulting in one-to-one competition, which does not outweigh conceptual priming. We use the swinging lexical network proposal as an instantiation of theories assuming both competitive lexical selection and conceptual facilitation to illustrate the predictions for the current experiment. However, please note that in the absence of explicit simulations, the net outcome of the trade-off between facilitation and competition predicted by the SLN in any particular situation remains somewhat vague. Thus, the intuitions we describe below should not be seen as specific to the SNL but rather more generally as consistent with the assumption that both competitive lexical selection and conceptual facilitation play a role during language production (Hantsch & Maedebach, 2013; Jescheniak, Matushanskaya, Mädebach, & Müller, 2014; Jescheniak, Schriefers, & Lemhöfer, 2014; Levelt, et al., 1999; Roelofs, 2018; Roelofs, Piai, & Schriefers, 2013a, 2013b; Starreveld, La Heij, & Verdonschot, 2013).

How would semantic richness be expected to modulate object naming from such a perspective? Many semantic features associated with the to be named concept should induce facilitatory effects similar to conceptual priming due to higher activation levels of concepts associated with many as compared to few semantic features (see Rabovsky & McRae, 2014, simulation 3, for evidence from a neural network model). This should result in increased activation flow to the corresponding lexical representation, which should induce faster lexical selection and naming. Indeed, in a recent behavioral study, we observed faster and more accurate naming for concepts associated with many semantic features (Rabovsky, Schad, & Abdel Rahman, 2016). The increased activation flow to the corresponding lexical representation may be accompanied by the activation of more co-activated lexical competitors – those sharing semantic features with the target. However, the lexical co-activation may not be strong enough to outweigh direct conceptual facilitation due to semantic feature activation. In contrast, a related variable, the density of semantic space, should be more directly related to lexical competition. This variable, called intercorrelational feature density, is also provided in the feature production norms by McRae et al. (2005). It indicates the degree to which a concept’s features are intercorrelated. Specifically, the authors constructed a matrix where each element corresponds to the production frequency of a specific feature for a specific concept, and then calculated pairwise correlations between the resulting feature vectors for all features that appeared in at least three concepts. The percentage of shared variance between each pair of a concept’s features (for feature pairs sharing at least 6,5% of their variance) was then summed. Concepts with high intercorrelational feature density inhabit denser regions of semantic space, and their activation entails stronger partial coactivation of other concepts through the intercorrelated features. Specifically, we assume that concepts with high intercorrelational feature density co-activate other mutually related concepts through the intercorrelated features. Highly correlated feature clusters often characterize groups of closely interrelated concepts (e.g. has wings, can fly, has a beak, can sing, etc.), and thus we assume that concepts with high intercorrelational feature density should co-activate a cohort of strongly interrelated lexical competitors. The activation of lexical cohorts should result in increased competition at the lexical level. This in turn is expected to induce substantial interference effects that override any possible facilitation brought about by conceptual coactivation. Indeed, in the behavioral study mentioned above, we observed object naming to be slower and more error-prone for concepts with high feature density (Rabovsky et al., 2016). This inhibitory influence of semantic feature density seems in line with studies investigating influences of semantic neighbors. Specifically, as noted above Mirman (2011) and Fieder, Wartenburger, and Abdel Rahman (2018) observed inhibitory influences of the number of closely related semantic neighbors (but see Bormann, 2011, for a null effect of overall semantic neighborhood size). Crucially, while this inhibitory influence is naturally predicted by language production theories assuming lexical competition, in the absence of external context manipulations semantic interference effects cannot directly be explained by non-competitive models.

What is the expected time course of these observed influences of item-endogenous semantic attributes? A comprehensive meta-analysis (Indefrey, 2011; updated based on previous version by Indefrey & Levelt, 2004; see also Strijkers & Costa, 2016, for discussion) suggests that lexical selection starts at around 200 ms poststimulus, when conceptual preparation has been completed, or at least, relevant semantic information to initiate lexical selection is available. Indefrey notes the possibility that “more specific conceptual information needed to select a particular lemma among a number of competitors” may come in later (p. 7) which is in line with the idea that conceptual and lexical processes interact and overlap in time (e.g., Abdel Rahman & Melinger, 2009b; 2019; Levelt et al., 1999). By now quite a few studies report an increased posterior positivity starting between 200 and 250 ms for situations presumably involving enhanced lexical competition such as cumulative semantic interference as discussed above (200 to 380 ms; Costa, Strijkers, Martin, & Thierry, 2009), especially for closely related category members with high feature overlap (250 to 400 ms; Rose & Abdel Rahman, 2016), as well as semantically related as compared to unrelated distractor words in a picture word interference paradigm (Rose et al., 2019; 200 to 300 ms). Because the inhibitory influence of feature density presumably also reflects lexical competition - even though not induced by a context but instead by the correlational structure of a concept’s features - this effect should be reflected in a similar enhanced posterior positivity starting between 200 and 250 ms.

How is the facilitatory influence of semantic richness expected to modulate ERPs? A similar time course would be in line with accounts functionally localizing semantic interference and facilitation at the same or interacting processing stages such as the model by Levelt et al. (1999; Roelofs, 2018) which assumes bi-directional links and continuous information transmission between semantic and lexical layers during speech production, or the swinging lexical network which similarly postulates conceptual facilitation and lexical competition to occur in parallel (Abdel Rahman & Melinger, 2009; 2019). On the other hand, theories such as the response exclusion hypothesis (Mahon et al., 2007) link facilitation and interference to distinct stages such that they would predict different temporal dynamics for both effects. As noted above, in line with lexical competition models we suggest that the feature density effect reflects lexical competition, which occurs in parallel with the conceptual facilitation induced by high semantic richness, such that we expect a similar time course for both influences. However, we had less clear expectations concerning the polarity of the semantic richness effect. On the one hand, based on the opposing influences of richness and density on naming times, one might expect opposing influences on ERP amplitudes as well. On the other hand, both a high number of semantic features and high feature density presumably increase activation in the lexico-semantic system (with the activation more specifically related to the to-be-named concept for high semantic richness, and more distributed across competing candidates for high feature density). This overall activation increase might be reflected in a similar enhanced posterior positivity for both, high richness and high density.

Summing up, we aimed to replicate previously observed facilitative influences of semantic richness and interfering influences of semantic density on object naming, and examine the temporal dynamics of these influences by means of ERPs. Please note that due to the focus on concept-inherent endogenous semantic features, in contrast to most studies on semantic context effects, the object names in the current study necessarily differ between experimental conditions, raising issues concerning potential confounding variables. In analogy to our behavioral study, we did not dichotomize the continuous variables, which can result in a loss of statistical power and the selection of unusual stimulus materials (Hauk, Davis, Ford, Pulvermuller, & Marslen-Wilson, 2006). Instead, we used 345 object concepts from McRae et al. (2005)’s norms, which were selected out of the overall 541 objects based on the sole criterion that they should have been named correctly by at least 75% of participants in our previous experiment (Rabovsky et al., 2016). Richness and density continuously varied in the stimulus set. Naming responses as well as ERPs were analyzed with linear mixed models (Baayen, Davidson, & Bates, 2008), which included potentially confounding variables (e.g., lexical frequency, visual complexity, etc.) as control covariates to allow for statistical control of confounds.

## 2 Material and methods

### 2.1 Participants

31 native German speakers (15 women) with mean age of 25 (range = 18 to 33) took part in our study. Six additional participants participated, but their data had to be excluded from analysis due to technical problems or excessive EEG artifacts. All participants reported normal or corrected-to-normal visual acuity, and all of them were right-handed according to a standard handedness questionnaire (Oldfield, 1971). All participants gave written informed consent prior to participation and received either course credit or monetary compensation (7€/ hour) for their participation. The study was approved by the ethics committee of the Psychology Department at Humboldt University Berlin.

### 2.2 Materials and procedure

Stimuli were grayscale photographs of 345 concrete object concepts from the feature production norms by McRae et al. (2005), where more than 700 participants were asked to list semantic features for 541 concepts and all features were retained that were produced by at least five of 30 participants. All pictures were scaled to 3.5 x 3.5 cm and presented on a light blue background. The 345 out of the overall 541 objects concepts from the norms by McRae et al. (2005) were selected based on previous naming performance, with only those object pictures included in the current stimulus set that were named correctly by at least 75% of participants in our behavioral study (Rabovsky et al., 2016). We used one-tailed Welch’s t-tests for unequal variances to compare the complete stimulus set of 541 concepts with the reduced set of 345 concepts concerning feature density and semantic richness. Feature density tended to be lower (*M* = 156.4, *SD* = 186.43 vs. *M* = 175.1, *SD* = 205.85; *t* = −1.4, *p* = .081) and semantic richness tended to be higher (*M* = 13.76, *SD* = 3.58 vs. *M* = 13.41, *SD* = 3.52; *t* = 1.43, *p* = .076) in the reduced stimulus set. There was a positive correlation between semantic richness and intercorrelational feature density in the reduced stimulus set, *r* = .46, *t*(343) = 9.63, *p* < .0001 (very similar to the correlation of *r* = .48 in the original stimulus set; Rabovsky et al., 2016). Participants were instructed to name the pictures as specific and accurate as possible. Each trial began with a fixation cross displayed in the center of a light blue screen for 0.5s. Then a picture was presented until a response was given or for a maximum of 4s. Naming latencies were registered with a voice key. After each trial, the experiment paused and the experimenter coded response accuracy using four categories: (1) experimental error (e.g. a noise made by the participant which was registered by the voice key) or time-out, (2) wrong naming response, (3) almost correct naming response, e.g. a synonym, or (4) correct (i.e., pre-determined) naming response. Once accuracy was coded, the experimenter initiated the continuation of the experiment; this procedure caused variation of the inter-trial interval (ITI). The 345 pictures were presented in a different random order for each participant, and the complete stimulus set was presented twice, resulting in a total of 690 trials, subdivided into six blocks of 115 objects each, which were separated by short breaks (blocks 4-6 were repetition blocks).

### 2.3 EEG Recording and Analysis

The continuous electroencephalogram (EEG) was recorded with Ag/ AgCl electrodes from 62 sites according to the extended 10-20 system, and referenced to the left mastoid. The vertical and the horizontal electrooculogram (EOG) were measured with peripheral electrodes below and left to the left eye. Electrode impedances were kept below 5 kΩ (below 10 kΩ for peripheral sites). Bandpass of amplifiers (Brainamps) was 0.032 Hz to 70 Hz, and sampling rate was 500 Hz. Offline, the EEG data were re-referenced to an average reference which has been recommended as being less biased than other common references (Picton et al., 2000). Then, a low-pass filter with a cut-off frequency of 30 Hz (24 dB/oct) was applied. Eye movements and blinks were corrected based on a spatio-temporal dipole modeling procedure using BESA software (Berg and Scherg, 1991) based on data from a short session recording prototypical individual eye movements and blinks at the end of the experiment. The continuous EEG was segmented into epochs of 1100 ms: 100 ms pre-stimulus onset (which served as baseline) until 1000 ms post-stimulus onset. Segments with missing or incorrect responses or artifacts were excluded. Artifacts were defined as potentials exceeding 50 *μ* V voltage steps between subsequent sampling points, a difference larger than 200 *μ* V within a segment, or amplitudes smaller than −200 *μ* V or larger than 200 *μ* V. Analyses focused on a cluster of posterior electrode sites comprising CP3, CP4, P3, P4, P5, P6, PO3, PO4, POz (based on Costa et al., 2009). For this region of interest (ROI) we conducted analyses using (generalized) linear mixed models (GLMMs) on mean amplitudes in a time segment selected based on previous findings concerning the posterior positivity in language production research, i.e. 200 to 550 ms (Aristei et al., 2011) to test for influences of semantic richness and semantic density as well as their interaction with repetition. Subsequently, we analyzed consecutive 10 ms segments between 0 and 1000 ms poststimulus within the same electrode cluster to explore the temporal dynamics of the observed effects in detail. The materials, data, and code for this study are available on the Open Science Framework (Rabovsky, Schad, & Abdel Rahman, 2019, https://osf.io/8wtp6/).

## 3 Results

We used (generalized) linear mixed models (GLMMs) implemented in the lme4-package (Bates, Maechler, Bolker, & Walker, 2014) in the R-system for statistical computing (www.r-project.org) to analyze influences of the number of semantic features and feature density on response times (recoded as 1/RT [sec] to adhere to normal distribution assumptions, via a LMM) as well as ERP amplitudes (for which we visually confirmed the approximation to normal distribution). Fixed effects were mean-centered for analysis. We z-transformed covariates to ease model fitting, and report standardized regression coefficients (β) to facilitate comparison between predictors, in addition to unstandardized regression coefficients (b) and the corresponding standard errors (SE). We also included repetition as a factor (using effect coding: first presentation = −0.5/ repetition = + 0.5), as well as its interactions with the number of semantic features and feature density. Moreover, we added crossed random effects for subjects and items, with random subject intercepts and uncorrelated random subject slopes for the repetition factor, the number of semantic features and feature density, and for response times their interaction with repetition. In addition, we controlled for influences of familiarity, number of orthographic neighbors, lexical frequency, as well as subjective (based on a rating) and objective (based on compressed file size) visual complexity and their interaction with repetition, without adding random slopes for these control covariates as suggested by Barr, Levy, Scheepers, and Tily (2013). For analyzing quadratic effects, we use the R-function *poly* in the *stats* package. For the LMMs, statistical testing was based on the Satterthwaite approximation for the denominator degrees of freedom as implemented in the lmerTest package (Kuznetsova, Brockhoff, & Christensen, 2014). For errors, logistic GLMMs using the *glmer* function in the *lme4* package failed to converge. We therefore instead used a Bayesian analysis with the same fixed-effects and a maximal random effects structure. We fitted this using the R function *brm* in the *brms* package (Bürkner, 2017; Bürkner, 2018), which estimates (e.g., hierarchical linear) Bayesian models in the probabilistic programming language *Stan* (Stan Development Team, 2017) using Hamiltonian Monte Carlo simulations of the posterior. We report estimates with 95% credibility intervals (CrI), and perform hypothesis testing using the R function *hypothesis* in the *brms* package. For the reported analyses, responses were considered correct when they received a score of 3 (almost correct, e.g. a synonym) or 4 (correct, i.e. pre-determined response) in the above described categories, analogous to our behavioral study (Rabovsky et al., 2016).

### 3.1 Performance

Response times as a function of the number of semantic features and intercorrelational feature density are displayed in Fig. 1; accuracy data are shown in Fig. 2. Object naming was more accurate for concepts with many semantic features (*β* = .37, *b* = 0.10, credibility interval, *95% CrI* [0.017, 0.19], *p*(b>0) = .989), and more error-prone for concepts with high feature density (*β* = −.39, *b* = −0.0022, *CrI95%* [−0.0037, −0.0006], *p*(b<0) = .995). None of these semantic effects was influenced by repetition (semantic features x repetition: 95% *CrI* [−0.034, 0.028]; density x repetition: 95% *CrI* [−0.00040, 0.00068]; see also Suppl. Fig. S1). Furthermore, object naming was faster for concepts with many semantic features (*β* = 0.031; *b* = 0.0086, *SE* = 0.0022, *t*(324) = 3.84, *p* = .0001). There was no main effect of feature density on RTs (*β* = −0.0082, *b* = .000045, *SE* = 000043, *t*(336) = −1.04, *p* = .30), but there was an interaction between repetition and feature density (*β* = −0.0019, *b* = .000057, *SE* = 0.000021, *t*(348) = −2.68, *p* = .0077), with significantly slower naming for concepts with high density during the repetition (*β* = −0.012, *b* = −0.000068, *SE* = 0.000039, *t*(297) = −1.72, *p* = .044, one-tailed based on the results from our behavioral study; Rabovsky et al., 2016), but not during the first presentation (*β* = −0.0032, *b* = −0.000018, *SE* = 0.000049, *t*(334) = −.36, *p* = .72).^1^ As to be expected, naming was overall faster (*β* = 0.00010, *SE* = 0.00001, *t*(30) = 8.91, *p* < .0001) and more accurate (*β* = .34, *CI* 95% [.24, .34]) during the repetition as compared to the first presentation. Based on visual inspection of Figure 1, we also ran an additional analysis testing for potential (unpredicted) quadratic, i.e. U-shaped effects of feature density on RTs. The main quadratic effect of density was not significant, *b* = −1.52, *SE* = .95, *t*(314) = −1.59, *p* = .11. There was an interaction between the quadratic effect of density and repetition, *b* = 1.15, *SE* = .47, *t*(13310) = 2.43, *p* = .015, with follow up tests showing a trend for a quadratic density effect during the first presentation, b = −2.0, *SE* = 1.1, *t*(327) = −1.85, *p* = .065, and no quadratic density effect during the repetition, *b* = −1.1, *SE* = 0.9, *t*(330) = −1.21, *p* = .22. As the quadratic density effect during the first presentation was unpredicted and only marginally significant, we will not discuss it further.

**Figure 1.**
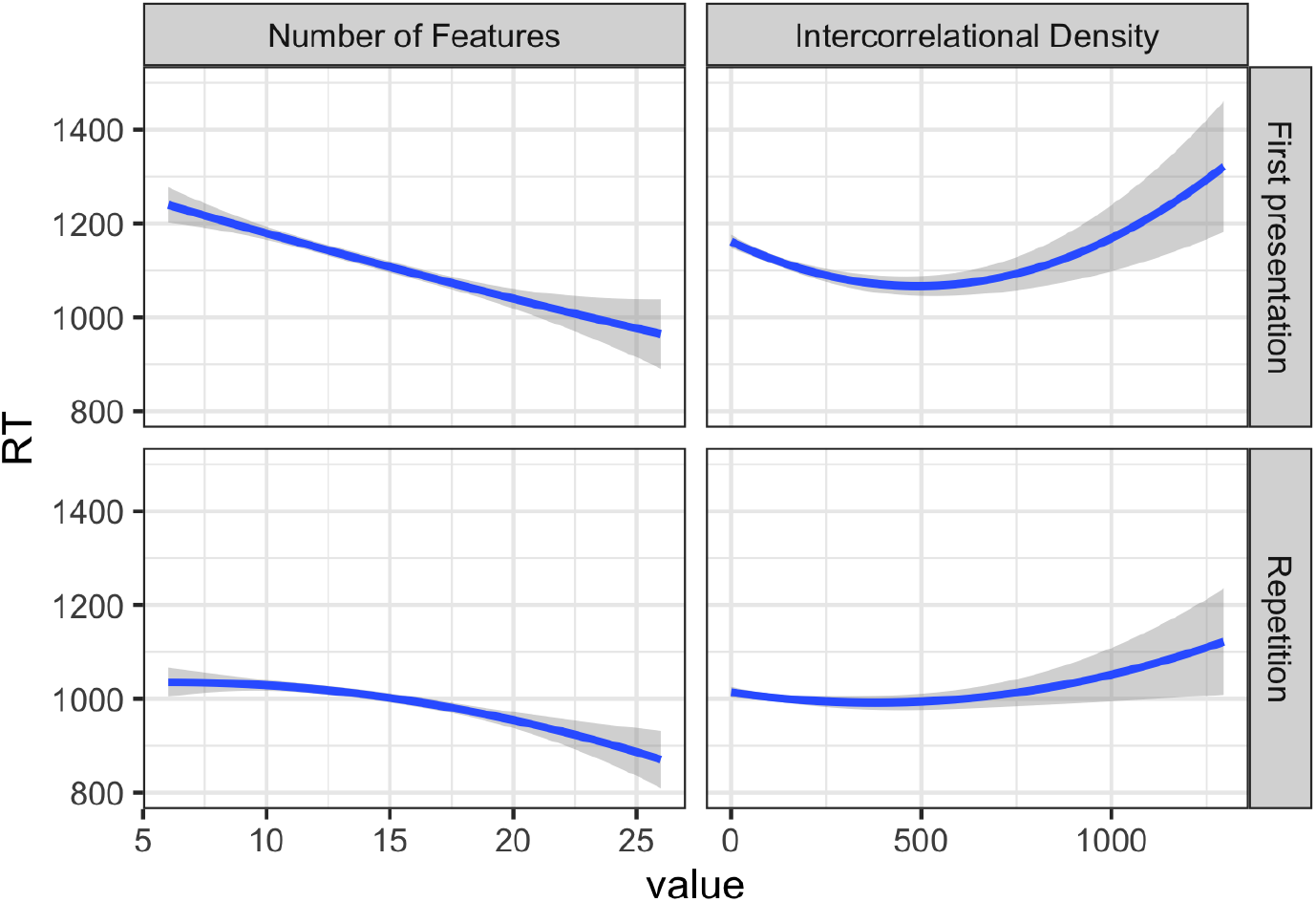
Response time (RT) as a function of the number of features (left) and feature density (right), during first presentation (top) and repetition (bottom) depicted as independently computed quadratic regression lines with 95% confidence bands.

**Figure 2.**
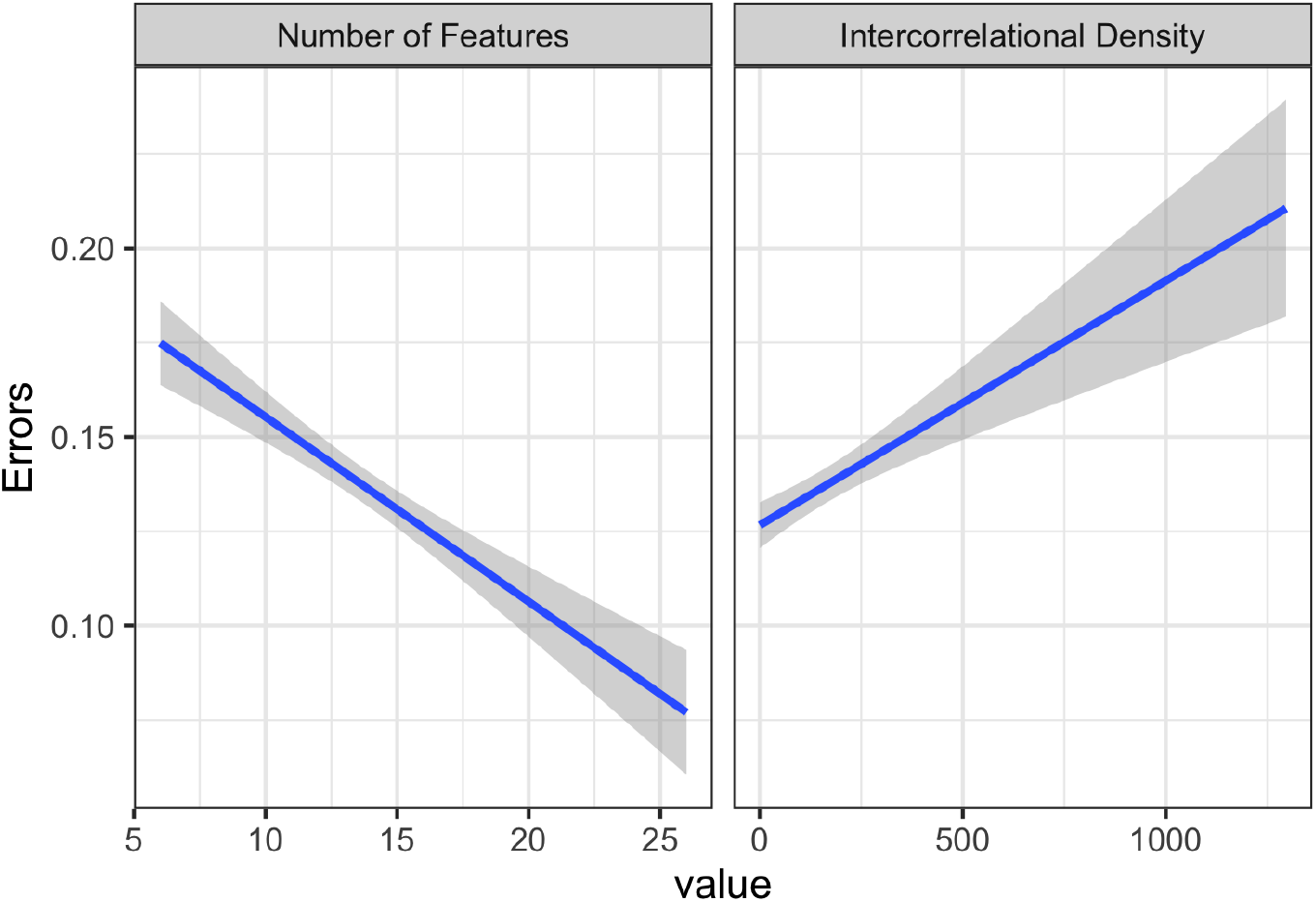
Accuracy as a function of the number of features (left) and feature density (right), depicted as independently computed logistic regression lines with 95% confidence bands.

### 3.2 Electrophysiology

ERP amplitudes at the posterior ROI as a function of semantic richness and feature density are depicted in Figure 3. It can be seen that there was an enhanced posterior positivity from about 250 ms onwards for both, concepts with many semantic features and concepts with high feature density. Indeed, both effects were significant in the pre-selected time segment between 200 and 550 ms, *β* = .48, *b* = 1.32, *SE* = .54, *t*(136) = 2.45, *p* = .015, for semantic richness, and *β* = .57, *b* = .031, *SE* = .011, *t*(122) = 2.86, *p* = .005 for feature density (see Fig. 3 for details on the time course of both effects). The density effect on ERPs was significant during more time segments during the repetition as compared to the first presentation (see Suppl. Fig. S2a for a depiction of ERPs plotted separately for first presentation and repetition), which might initially seem as if the increase of the density effect observed in the behavioral data was mirrored in ERPs. However, the interaction between repetition and both, feature density and the number of features, did not reach significance, neither in the average segment between 200 and 550 ms, *t*s < |.84|, *p*s > .4, nor in any of the 10 ms segments within this interval (see Suppl. Fig. S2b for a depiction of the interaction between repetition and the semantic variables).

**Figure 3.**
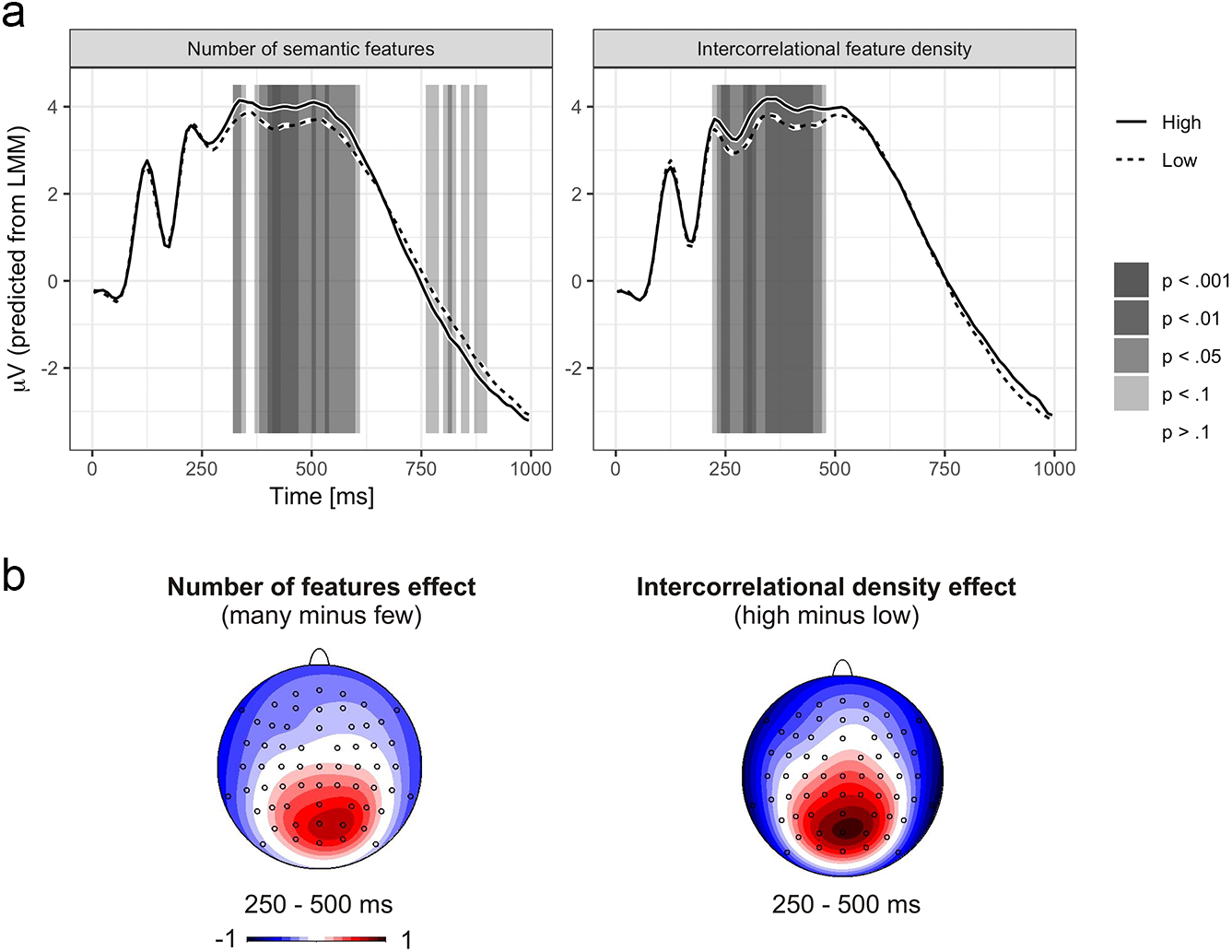
a. Depicted are linear mixed model estimates of the mean voltages plus/ minus the effect sizes (resulting in depictions of waveforms for high/ low conditions) of the influences of the number of semantic features (left) and feature density (right) at a posterior region of interest (CP3, CP4, P5, P3, Pz, P4, P6, Po3, POz, PO4; based on Costa et al., 2009) in consecutive 10 ms segments between 0 and 1000 ms. Grey shading indicates levels of significance. b. Topographical distributions of the effects are based on median-splits.

Figure 4 displays the time course of the correlation between ERP amplitudes at the posterior ROI and naming times over items. It shows that more positive amplitudes at this posterior ROI were related to slower naming times between 230 and 380 ms. Even though the correlation seems to be stronger and seems to last longer during the repetition (see Suppl. Fig. S3), the difference in the correlation during the relevant time segment between the first and second presentation was not significant (see Suppl. Fig. S4).

**Figure 4.**
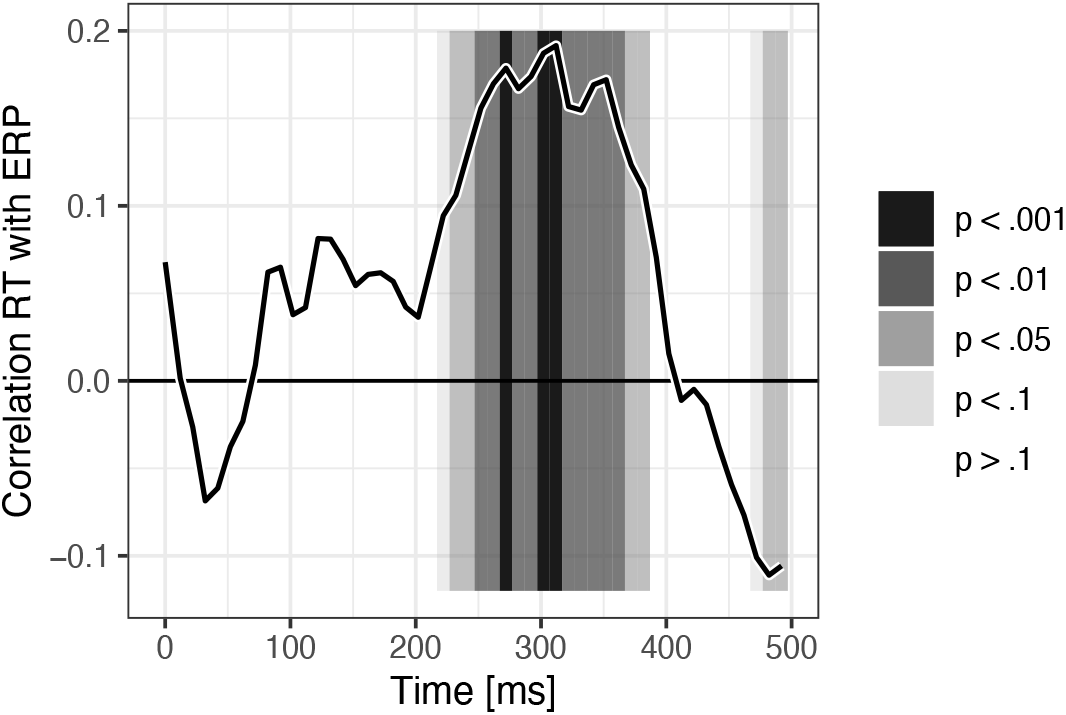
Correlation between naming times and ERP amplitudes at the posterior region of interest, averaged across items at both presentations.

## 4 Discussion

The present study demonstrates that both, facilitating influences of semantic richness, and inhibitory influences of semantic density (based on McRae et al., 2005), are reflected in an enhanced posterior positivity in the ERP starting at about 250 ms. While naming times and accuracy reflect facilitated production of semantically rich messages that are associated with many semantic features, naming is slower (during the second presentation) and more error-prone for concepts with high intercorrelational feature density (see Fig. 1 & 2). Both a high number of semantic features and high intercorrelational feature density induced an increased posterior positivity starting at about 250 ms (see Fig. 3), and amplitudes at the same posterior electrodes correlated positively with naming times between about 230 and 380 ms (see Fig. 4).

As discussed in the introduction, many semantic features presumably increase activation at the semantic level (see Rabovsky & McRae, 2014; simulation 3), enhancing the flow of activation to the corresponding lexical representation and thereby facilitating the naming response. On the other hand, possible inhibitory consequences of the activation of many semantic features that might be shared with other related concepts and their lexical representations may not be strong enough to override this facilitation. Instead, the co-activation of competing lexical candidates should be more directly induced by a related variable, the semantic feature density. The swinging lexical network proposal (SLN; Abdel Rahman & Melinger, 2009b) assumes that inhibitory consequences of lexical competition will outweigh facilitation at the conceptual level when an interrelated lexical cohort of sufficient size is activated. In the case of feature density, the co-activation of the semantic representations of other related concepts might partially feed back to the target concept, inducing some facilitation at the conceptual level. However, as highly correlated feature clusters typically characterize groups of interrelated concepts (e.g. has four legs, has fur, has a tail,…), items with high feature density should activate lexical cohorts that mutually increase their activation, resulting in one-to-many competition and therefore substantial interference which outweighs facilitative influences at the semantic level. As noted in the introduction, in the absence of explicit computational modeling, predictions of the net outcome of the interplay between competition and facilitation in SLN remain necessarily vague. However, in general, language production theories assuming both conceptual facilitation and competitive lexical selection can naturally account for the obtained pattern of facilitatory and inhibitory influences of the number of semantic features and their density in semantic space. On the other hand, it is not clear how the present findings would be explained by language production theories dismissing lexical competition. The response exclusion account by Mahon and colleagues (2009) was specifically proposed to cover distractor word effects in the PWI paradigm and is therefore not directly relevant for the present findings because no distractor or context stimuli are presented. Likewise, the model proposed by Oppenheim and colleagues (2010) offers an account that includes incremental learning as part of language production. This model can explain semantic blocking and cumulative interference effects, and it is not in conflict with our findings, but it cannot directly account for our message-endogeneous semantic effects.

The observed temporal dynamics with influences of both effects starting at about 250 ms are in line with previous evidence for electrophysiological indicators of lexical competition, e.g., in cumulative semantic interference (Costa et al., 2009; Rose & Abdel Rahman, 2016) as well as picture word interference and semantic blocking (Aristei et al., 2011; Rose et al., 2019). The time course is also roughly in line with the meta-analysis by Indefrey (2011) suggesting lexical competition to start at about 200 ms. The fact that the effects observed here start towards the end of the expected starting time might be due to contextual influences on lexical competition starting slightly earlier than endogenous influences on lexical competition: the context is activated before the item and thus already influences competition at the very beginning of lexical selection, while endogenous properties might depend on the item’s activation, and activation flow from the item to other related items, to unfold. This explanation would also be in line with the observation that the correlation between ERP amplitudes and naming times seems to start slightly earlier than the influences of the variables of interest, suggesting that lexical selection might have been influenced by additional factors with a slightly earlier impact.

Both ERP effects as well as the correlation of ERPs with RTs continued longer than the duration of lemma retrieval estimated by Indefrey (2011), which was 75 ms (Table 2 in Indefrey, 2011). However, it is important to note that the estimated articulation time in Indefrey’s time line is 600 ms which is considerably earlier than the average naming time of over 1000 ms observed in our study, and Indefrey himself notes that the different stages may vary considerably in length. Many object naming studies rely on relatively small sets of highly typical items (often simple line drawings) that are repeated many times, while we used a broad range of complex photographs of 345 different objects (e.g., robin or gown) which might have increased difficulty and thus time demands of many of the involved processing stages. Furthermore, as discussed in the introduction, Indefrey (2011) assumes that lexical selection starts as soon as relevant semantic information is available (e.g., animacy information), but additional semantic information may come in later to more specifically single out the to-be-produced lemma among competing candidates (e.g., among various types of birds or trees). Thus, conceptual processing may continue after lexical selection has been initiated, with both processes occurring simultaneously and in interaction as suggested by lexical competition models (Abdel Rahman & Melinger, 2009b; Levelt et al.,1999; Roelofs, 2018). This assumption is also supported by the simultaneous ERP effects of facilitating influences of semantic richness effects and of inhibitory influences of semantic density.

Even though the influences of semantic richness and feature density are in opposite direction in performance, ERP effects of both variables are of the same polarity. Specifically, high values of both semantic richness and feature density induced an increased posterior positivity. The functional basis of this posterior positivity seems not entirely clear. It seems to reflect aspects of lexical selection, presumably the difficulty of lexical selection, as indicated by the positive correlation with naming times. An important factor in the difficulty of lexical selection is lexical competition such that a likely candidate process underlying the observed positivity might be lexical competition. This would be in line with previous reports of similar positivities in paradigms inducing lexical competition such as cumulative semantic interference (Costa et al., 2009; Rose & Abdel Rahman, 2016) and picture word interference. It would also seem plausible in light of the interpretation of the density effect as inducing enhanced lexical competition. On the other hand, the observation that high semantic richness induced a similar enhanced positivity even though it entailed facilitated naming suggests that the story might be somewhat more complex. It seems possible that the enhanced posterior positivity reflects a less specific process than lexical competition, a process that covaries with lexical competition but is not limited to it, such as for instance activation in the lexical semantic system during competitive lexical selection. This activation might be high for both, high semantic richness and high feature density. It might be primarily related to the to-be-named concept for concepts with a high number of semantic features, and more distributed across competing candidates for concepts with high feature density. In both cases the overlapping time course and same polarity seems to suggest a high degree of overlap and possible interaction between conceptual and lexical processing (for a related discussion see Abdel Rahman and Melinger, 2019)

An interesting issue is how the posterior positivity observed during language production relates to the N400 ERP component reflecting semantic processing during comprehension (see Kutas & Federmeier, 2011, for review, and Rabovsky, Hansen, & McClelland, 2018, for an implemented computational model). The topographical distribution of both effects is similar, and in many situations, they would be expected to be modulated in the same way. For instance, the presentation of semantically related items, a semantically homogeneous composition of objects, and the previous processing of items from the same semantic category would all be expected to reduce the amplitude of the N400 component during comprehension. This would result in increased positivity, which corresponds to the pattern observed during production. However, during language comprehension, a decreased N400 and thus increased positivity is typically associated with facilitated processing. In contrast, during language production, increased positivity usually goes along with increased interference (see e.g., Blackford et al., 2012, for a language production study interpreting the observed ERP effect as an N400). In this context, it seems interesting to note that we investigated influences of the number of associated semantic features during visual word processing. In this study, we observed increased N400 amplitudes (and thus decreased positivity) for words with many features (Rabovsky, Sommer, & Abdel Rahman, 2012a; see also Rabovsky et al., 2012b; 2012c), in line with increased N400 amplitudes for concrete as compared to abstract words (Kounios & Holcomb, 1994). Thus, the polarity of the influence that we observed during comprehension differs from the effect observed in the current study for object naming. This suggests that the observed positivity may indeed reflect processes specific to language production, such as conceptually driven lexical selection, which is not required in comprehension tasks. Further research seems required to better understand the relationship between both types of ERP effects.

Another aspect of the data that seems worth discussing is the comparison of the influence of feature density during the first presentation and the repetition (Fig. 1), showing significantly slower naming times for concepts with high feature density only during the repetition while accuracy was significantly lower for high density concepts during both presentations (Fig. 2)^2^. This increase of the density effect with repetition seems plausible: If concepts have already been partially co-activated through intercorrelated features during the first presentation, this previous co-activation might enhance the concepts’ readiness to get co-activated more strongly during the repetition. Furthermore, beyond the repetition of the individual items, the intercorrelated features presumably occur in different categorically related items which all result in partial co-activation of the corresponding concepts. Some of the co-activated concepts might be the to-be-named concepts in other trials such that overall, the readiness for co-activation and the resulting lexical competition should be increased during the repetition. This seems to naturally explain the stronger influence of feature density during the repetition as compared to the first presentation within the frame of theories assuming competitive lexical selection (Abdel Rahman & Melinger, 2009a, 2009b; Levelt, et al., 1999; Roelofs, 2018).

In conclusion, the present study focused on investigating the electrophysiological correlates and comparing the temporal dynamics of influences of message-endogeneous attributes of semantic richness and density. While endogenous semantic influences on language production have recently attracted growing attention (Bormann, 2011; Fieder et al., 2018; Mirman, 2011; Rabovsky et al., 2016), their neural correlates and time courses in neurotypical subjects have not yet been investigated (for clinical evidence, see Fieder et al., 2016 and Mirman & Graziano, 2013). We replicated facilitative influences of semantic richness, operationalized as the number of features associated with a to-be-named object concept, and inhibitory influences for concepts that inhabit denser regions in semantic space, operationalized as intercorrelational feature density (Rabovsky et al., 2016). Both effects were reflected in an increased posterior positivity in the ERP starting at about 250 ms, which has been related to lexical selection and competition in previous work (e.g., Costa et al., 2009; Rose & Abdel Rahman, 2016). These findings demonstrate that the richness and density of semantic representations induce similar processes as those evoked by more common contextual manipulations in object naming, and play an important role in lexical selection during speech planning.

## Acknowledgments

This research was supported by the German Research Council (DFG) via grant AB 277/ 4-2 to Rasha Abdel Rahman and grant RA 2715/2-1 to Milena Rabovsky. We thank Yumi Tatsumiya and David Schröter for help with stimulus preparation and data acquisition and Guido Kiecker for technical assistance.

## Author note

Data and materials are made available on the Open Science Framework (OSF): https://osf.io/8wtp6/

## Footnotes

1 In addition to these primary analyses, we repeated RT analyses using bootstrapping and computing 95% bootstrap confidence intervals (based on 1000 iterations). The effects were again significant, including the number of feature effect (95% bootstrap CIs [0.015 0.047]), the interaction between the semantic density effect and repetition (95% bootstrap CI [0.018 −0.003]), and the semantic density effect during the second presentation (95% bootstrap CI [−∞ −.0003], one-tailed based on the results from our behavioral study, Rabovsky et al., 2016). We performed additional analyses, where raw RTs were analyzed with repeated measures GLMs not assuming a normal distribution, but rather an inverse Gaussian distribution, which fits the raw RTs well (see also Lo & Andrews, 2015). This analysis again confirmed the general pattern of RT results. Specifically, while the model failed to converge for 4 subjects, with 27 valid subjects there was still a significant facilitating effect of the number of semantic features, *t*(26) = 11,76, *p* < .0001, and a trend for an interaction between repetition and feature density, *t*(26) = −1.96, *p* = .061. For the analysis of RTs during the second presentation, there were 29 valid subjects (analysis failed to converge for 2 subjects) and there was a significant inhibitory effect of feature density, *t*(28) = −1.90, *p* = .034 (one-tailed based on Rabovsky et al., 2016).

2 A possible explanation for the difference with respect to our previous behavioral study where we observed an inhibitory influence of density already during the first (and only) presentation (Rabovsky et al., 2016) lies in the difference between the stimulus sets, with on average lower density values in the current reduced as compared to the previously used complete stimulus set (see section ‘Materials and procedure’).

## Notes

### Competing Interest Statement

The authors have declared no competing interest.

