## Supplementary material (Fig. S1-S4) for "Semantic richness and density effects on language production: Electrophysiological and behavioral evidence"

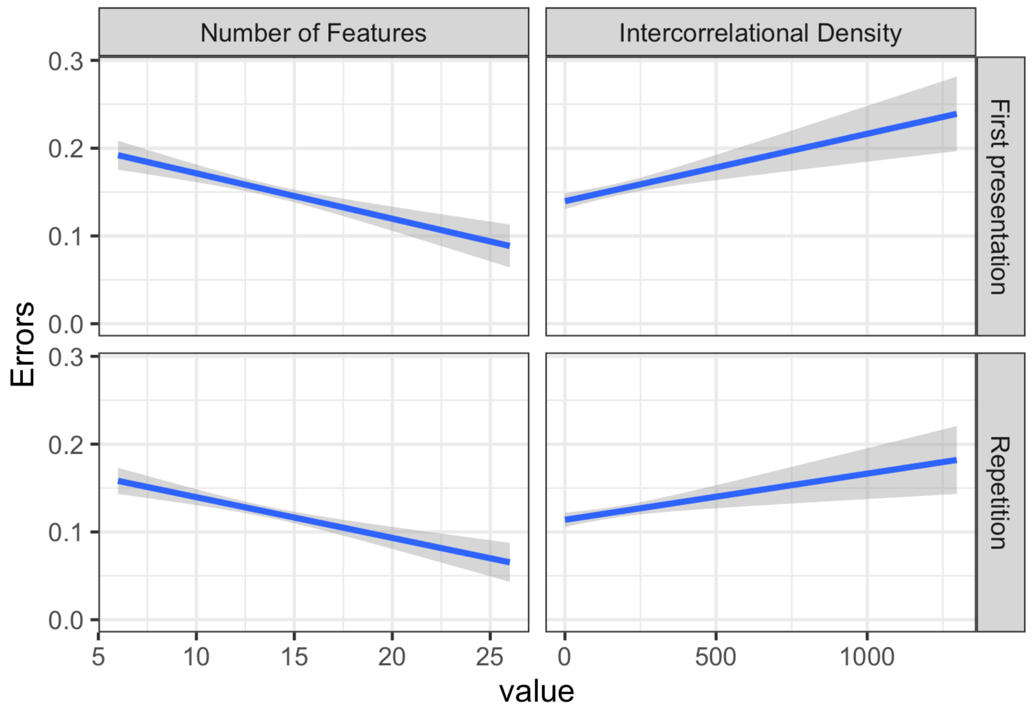


*Supplementary Figure S1. Accuracy as a function of the number of features (left) and feature density (right), during first presentation (top) and repetition (bottom), depicted as independently computed logistic regression lines with 95% confidence bands.*

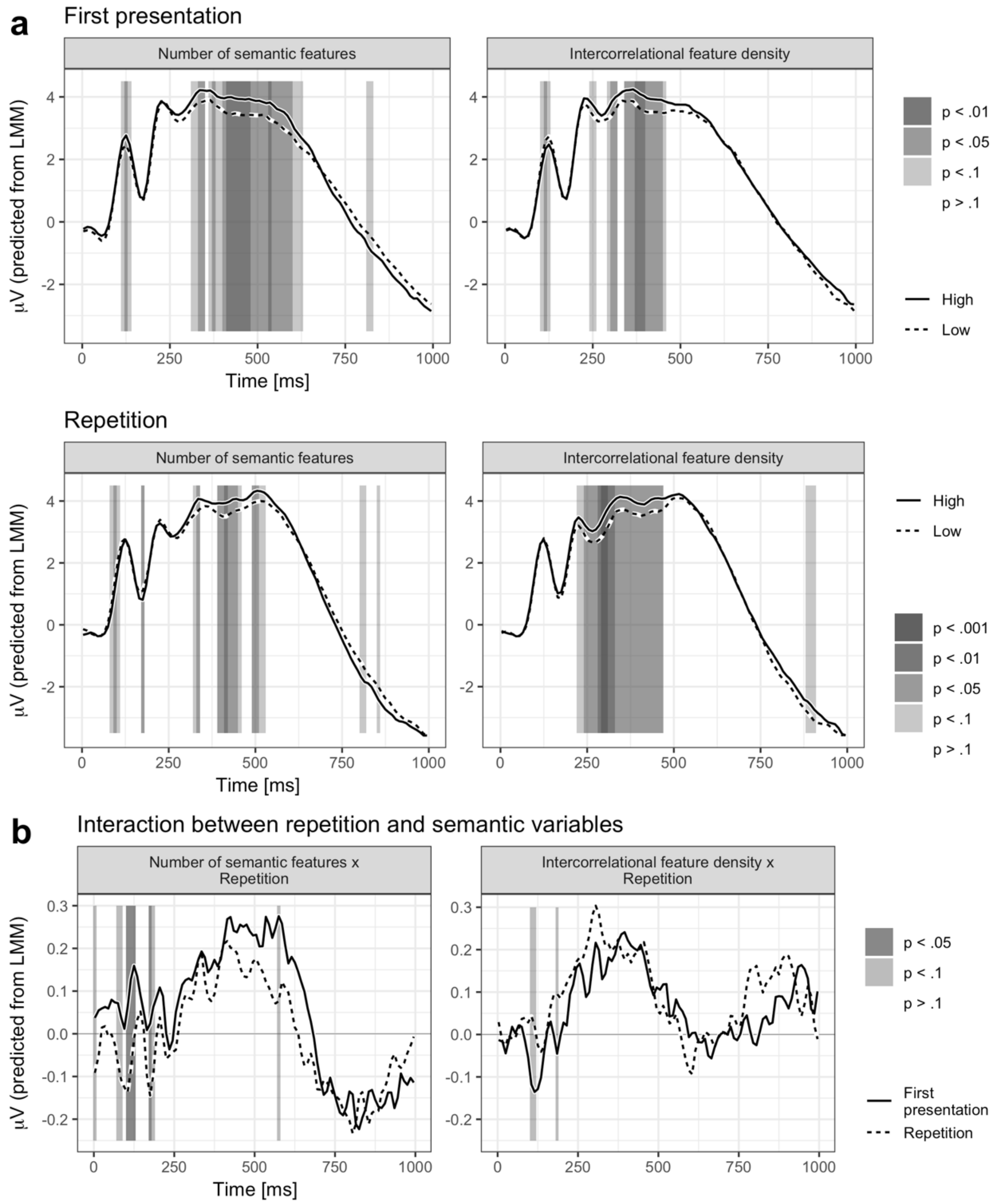


*Supplementary Figure S2. a. Depicted are linear mixed model estimates of the mean voltages plus/ minus the effect sizes (resulting in depictions of waveforms for high/ low conditions) of the influences of the number of semantic features (left) and feature density (right) during the first presentation (top) and the repetition (below) at a posterior region of interest (CP3, CP4, P5, P3, Pz, P4, P6, Po3, POz, PO4; based on Costa et al., 2009) in consecutive 10 ms segments between 0 and 1000 ms. Grey shading indicates levels of significance. b. Difference waves for influences of the number of features (many minus few; left) and feature density (high minus low; right) as a function of presentation condition (first presentation versus repetition). Grey shading indicates levels of significance of the interaction between repetition and the semantic variables.*

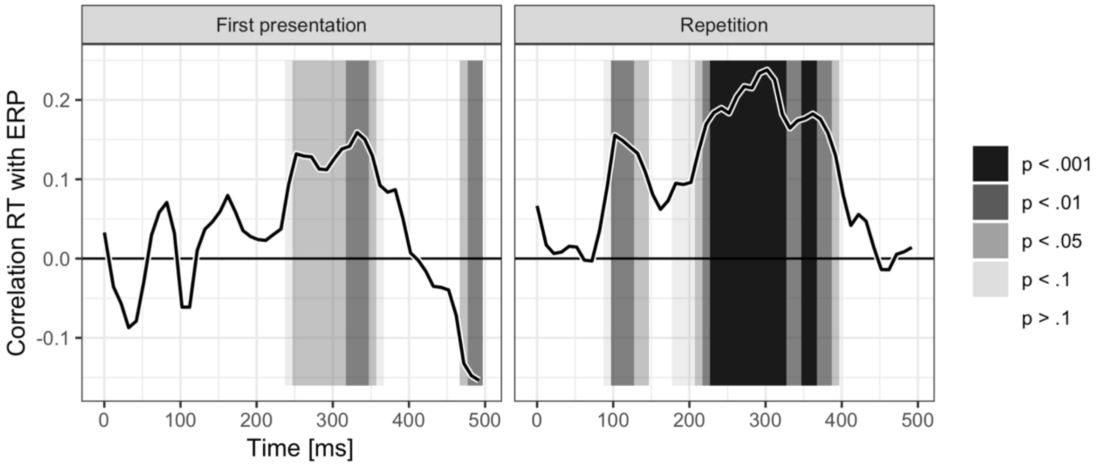


*Supplementary Figure S3. Correlation between naming times and ERP amplitudes at the posterior region of interest over items during the first presentation (left) and the repetition (right).*

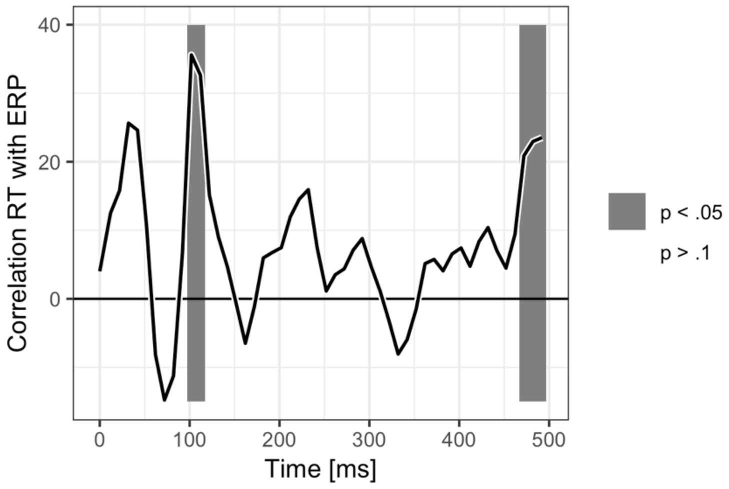


*Supplementary Figure S4. Repetition effect on the correlation between naming times and ERP amplitudes at the posterior region of interest over items.*
